# Egocentric Bias in Effort Comparison Tasks Is Driven by Sensory Asymmetries, Not Attribution Bias

**DOI:** 10.1101/2022.08.12.503607

**Authors:** Caedyn Stinson, Igor Kagan, Arezoo Pooresmaeili

## Abstract

When comparing themselves with others, people often perceive their own actions and behaviour favourably. This phenomenon is often categorised as a bias of attribution, with favourable self-evaluation resulting from differing explanations of one’s own behaviour and that of others. However, studies on availability biases offer an alternative explanation, ascribing egocentric biases to the inherent sensory asymmetries between performing an action and merely observing it. In this study, we used a paradigm that allowed us to directly compare these two distinct sources of bias. Participants perceived the tasks they performed to be harder than the tasks they observed, but demonstrated no bias driven by favourable self-evaluation. Furthermore, the degree of overestimation of the difficulty of performed tasks was magnified as overall task difficulty increased. These findings suggest that egocentric biases are in part derived from sensory asymmetries inherent to the first-person perspective.

## 1. Introduction

### 1.1 The Role of Social Comparison

Social comparison is performed in a plethora of cognitive processes. It is particularly evident when people consider both sides of an effort–reward trade-off: The value a person gives to a reward depends not just on the reward’s absolute value but also its value relative to that received by their peers (Boyce et al., 2010; Card et al., 2012; Fliessbach et al., 2007). People also monitor relative effort across their social group in order to identify social loafing (Karau & Williams, 1993) and to ensure that reward is meted out proportionately to effort (Akerlof & Yellen, 1990; Bland et al., 2017). Social comparisons also come into play in cognitive processes that require the integration of social information, such as using social peers as reference points to evaluate one’s own abilities (Dijkstra et al., 2008; Wolff et al., 2018), considering and evaluating opinions (Buunk & Gibbons, 2007; Festinger, 1954), incorporating the actions of others into the learning process (Dijkstra et al., 2008; Mattar & Gribble, 2005), and conforming to social norms (Allcott, 2011; Yun & Silk, 2011).

Comparisons between oneself and others are subject to a range of biases. In many situations, individuals struggle to objectively compare others’ behaviour to their own. Typically, these biases err in the direction of favourable self-assessment and as such are widely referred to as *egocentric* or *self-serving biases* (Blaine & Crocker, 1993; for a review see Greenberg et al., 1982). Egocentric biases include the overestimation of one’s contribution to cooperative tasks (Corgnet & Gunia, 2010; Kruger & Dunning, 1999; Ross & Sicoly, 1979); the better-than-average effect, in which more than 50% of people consider themselves to be above average in a given task (Zell et al., 2020); the willingness of feuding parties to pursue economically irrational legal actions (Loewenstein et al., 1993); and overestimation of one’s own impact on outcomes (Berberian et al., 2012). Unfavourable biases in social comparison have also been demonstrated, with people displaying a tendency to compare upwards when evaluating social standing (Lup et al., 2015) and wages (Harris et al., 2008) that can lead to dissatisfaction with one’s own situation.

#### 1.1.1 Explanations for Social Comparison Biases

Biases in social comparison have been extensively studied in the field of attribution theory (Weiner, 1985; Weiner et al., 1987). Attribution theory posits that biases in social comparison arise from asymmetrical attribution in people’s causal explanations of behaviour. According to the theory, explanatory sources are asymmetrically attributed for self and other to either external (i.e., situational) or internal (i.e., stemming from the actor) causes (Gilbert & Malone, 1995; Jones & Nisbett, 1987; Pronin, 2008). An important meta-analysis (Malle, 2006) found that people were more likely to attribute their own success to internal factors and their failures to external factors.

While these studies provide a wealth of evidence of social comparison asymmetries, they offer little in the way of causal explanation of the underlying mechanisms. Studies that investigate the information available to individuals, however, offer more insight into the matter. These studies suggest that asymmetries of attribution may be rooted in the fact that a person’s own intentions and the direct sensory input from their actions are simply more available to them than are the intentions and sensory input of others. This availability-driven bias is evident in the overestimation of one’s own contributions to group tasks (Ross & Sicoly, 1979), the underestimation of collaborators in group tasks with increased physical distance (Corgnet & Gunia, 2010), and the greater saliency of external factors that impede people from attaining their goals compared to factors that assist them (Davidai & Gilovich, 2016). These findings suggest that when comparing one’s own actions with those of someone else, information about one’s own experience is accessed more readily than is information about others’ experience, and that this phenomenon drives the observed egocentric bias.

Crucially, however, these studies do not distinguish the psychophysiological origin of this biased perception. Specifically, while the explanations are typically framed as an asymmetry between *self* and *other* (e.g., Pronin, 2008; Weiner, 1985), or *active* and *observe* (e.g., Jones & Nisbett, 1987; Malle, 2006), the paradigms used in studies are insufficient to distinguish between these two potential sources of asymmetry. This distinction between sources of asymmetry is important. Biases driven by self/other asymmetries imply that sensory inputs of personal experiences are noisy but unbiased, and that bias originates from favourable representations of oneself over others. Biases driven by active/observe asymmetries imply that the source of bias is the asymmetry in sensory information inherent to the first-person perspective, with internal representations and the subsequent recall of experiences being noisy but unbiased.

For example, an individual exhibiting a self/other bias would unbiasedly receive information related to both their own and their partner’s contribution to household tasks—but their preconceived representations (e.g., “ my partner is typically lazy” /” I normally get housework done quickly”) would lead them to a biased interpretation of the information. An individual exhibiting an active/observe bias would receive more information about their own contribution than about their partner’s contribution. This asymmetry of information would lead to the biased perception that they contribute more to the housework. These differences may initially appear minor—after all, both individuals would be more likely to claim that they do more housework than their partner—but the implications are not. A self/other bias indicates a representational bias that has been reinforced over a lifespan (Palminteri et al., 2017) and is likely to resist change, whereas an active/observe bias is driven by information availability and therefore can be corrected swiftly by making information more symmetrical (e.g., through perspective taking; Richardson et al., 2021; Zhou et al., 2013).

We aimed to dissociate the two potential sources of egocentric bias—self/other asymmetry and active/observe asymmetry. To do so, we used an effortful task previously used to demonstrate that individuals incorporate both external (reward) and internal (sensory) information in a Bayes-optimal manner when assessing both their own (Pooresmaeili et al., 2015) and a partner’s effort (Rollwage et al., 2020). In these studies, the paradigm elicited asymmetry in the degree to which participants incorporated the reward information: Participants incorporated reward information to a lesser extent on their own trials than when they observed a partner. This result suggested that participants were also asymmetrically incorporating their sensory information, with their own effort playing a greater role in their decision than the effort they observed their partner making. We aimed to exploit this apparent asymmetry in sensory processing by removing the reward information and instead having participants directly compare the difficulty of the task when performed by themselves and a partner. By adding a condition in which participants also watched their own prerecorded performance of the task, we were able to distinguish between biases driven by self/other and active/observe asymmetries.

We hypothesized that we would detect both self/other and active/observe biases at the group level. We also predicted that participants would exhibit greater accuracy when assessing the difficulty of tasks they performed compared to those they merely observed, and that the accuracy disparity between *active* and *observe* conditions would correlate with the degree of active/observe bias, but not with self/other bias.

We found that participants perceived tasks they performed to be more difficult than tasks they observed, but found no significant bias between *self* and *other* conditions. We investigated whether differences in perceptual accuracy between the conditions were responsible for the observed bias, however, while participants judged the tasks they performed more accurately than those they observed, accuracy differences between *active* and *observe* conditions did not correlate with an active/observe bias. Instead, our exploratory analysis suggested that active/observe bias was driven by overall task difficulty.

## 2. Methods

### 2.1 Participants

The study was approved by the Freie Universität Berlin Department of Educational Science and Psychology Ethics Commission. A preregistered power analysis using data from Rollwage et al. (2020) indicated that a sample size of 49 was required. To account for participant dropout or exclusion, 63 participants were recruited, with 51 adults aged 18–35 years old included in the final analysis (31 female, 20 male; mean age 24.4; SD = 4.1 years; 45 right-handed). Of the 12 individuals who participated in the study but were excluded from analysis, 10 were excluded for not meeting the required accuracy threshold of >80% correct on trials where the easiest task was compared with the most difficult, one was excluded for correctly identifying the experimental manipulation in a post-experimental survey, and one was excluded because their performance fell 4.20 standard deviations from the sample mean.

Participants were recruited from Freie Universität Berlin via mailing lists and on-campus advertising and from social media groups for English speakers living in Berlin. Participants were required to have normal/corrected to normal vision, no current health problems, no history of psychiatric or neurological disorders (self-reported), and sufficient English skills to understand the task instructions and the consent documents. Participants received a base payment of €8/hour (the experiment took 4 hours) plus bonus payments to encourage performance and attention: Extra money was offered for completing *active* tasks, for two-interval forced choice (2IFC) accuracy, and for correctly detecting ball colour changes on selected trials (see section 2.2.1 for the description of this task). The maximum payment for the complete study was €45.34. Students of Freie Universität Berlin could opt to receive course credits in lieu of the base payment. To minimise inequality in participant incentives, these students received their overall winnings without the hourly pay, for a maximum of €13.34. Of the 55 participants who completed the full study (four of whom were excluded), 30 were fully paid, receiving a mean payment of €42.24, and 25 received four course credit hours and a mean payment of €8.94.

### 2.2 Apparatus and Stimuli

The experimental stimuli was produced using MATLAB (2016a, The MathWorks, Inc., Natick, Massachusetts, United States) and Psychophysics Toolbox (Brainard, 1997; Pelli, 1997). Four desktop PCs were used (6GB RAM; Intel® Core(TM) 2 Duo Processors E8400) with Samsung SyncMaster monitors (model: 943BR; screen width 29.4 cm; screen height 16.6 cm; resolution 1280×1024 pixels), and Dell keyboards (model: E145614). The screen was placed 50 cm from the estimated viewing position. The computers were positioned so that participants were unable to see the other participants’ screens.

The task featured a white ball (radius = 30 pixels), and a ramp that culminated in an upper plateau (ramp length = 550 pixels, angle = 30°, plateau length = 150 pixels; Figure 1) with a black background. Participants moved the ball up the ramp by pressing left and right keys in alternating order and were instructed to use their dominant hand for the entire experiment. Each key press resulted in a constant amount of displacement of 25 pixels up the ramp. Participants wore over-ear high ambient noise-attenuating headphones (Sennheiser HD 280/380 Pro), which delivered a beep for every key press. The ball’s progress was opposed by a “ gravity level” that rolled the ball down the ramp at a constant rate. If participants stopped pressing keys, or pressed too slowly, the ball would roll back toward the start point, where it would remain until key pressing resumed. A task consisted of either actively moving the ball (*active* condition) or watching the ball being moved (*observe* condition). Each 2IFC trial consisted of two tasks. After each trial participants rated the relative task difficulty of the two tasks using a rating bar (1024 pixels wide x 102 pixels high) and rating slider (10 pixels wide x 204 pixels tall).

**Figure 1.**
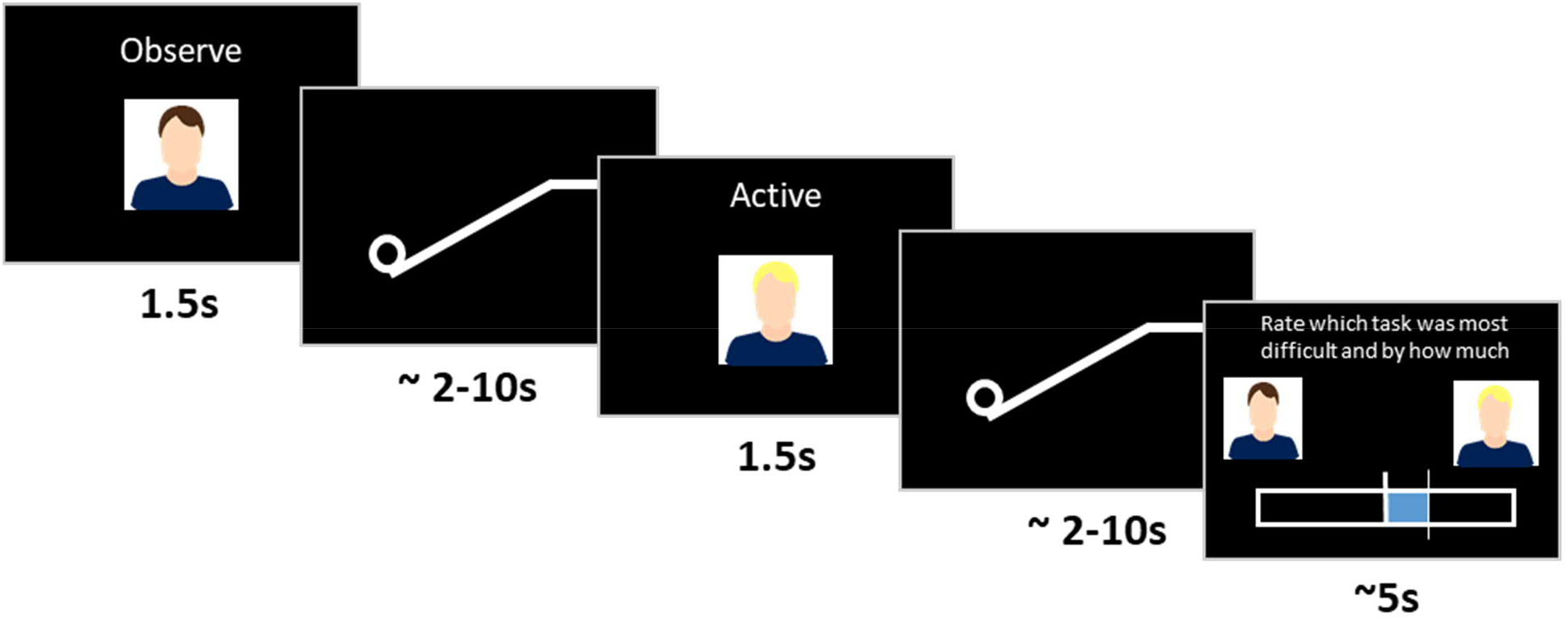
Two-Interval Forced Choice Workflow. *Note*. Each trial comprised two tasks (pushing a ball, or watching a ball be pushed up a ramp), performed by participants themselves (*self* condition) or someone else (*other* condition). Before each task, a photo of the person who performed the task and whether the participant had to perform the task (*active* condition) or observe it being performed (*observe* condition) was displayed onscreen. After the two tasks were completed, participants rated which was more difficult.

### 2.2 Experimental Design

#### 2.2.1 Paradigm Overview

The ball and ramp task had three conditions:

- *self*_*active*_ : Participants used the keyboard to move the ball up the ramp.
- *other*_*observe*_ : Participants watched a recording of what they were told was their partner moving the ball up the ramp (but was in fact a recording of their own performance).
- *self*_*observe*_ : Participants watched a recording of themselves moving the ball up the ramp.

Each trial consisted of two tasks presented in sequential pairs, with individuals performing a 2IFC in which they used a slider on a continuous rating bar to indicate which of the two tasks they thought was more difficult (i.e., which had the greater gravitational force) and by how much. Participants had to select one of the two items and were unable to register the tasks as equally difficult. This continuous rating schema provided both binary data (which task was more difficult) and continuous data (the first task was *x* more/less difficult than the second task). A cover task where the ball would briefly change colour on one of the two tasks was also included so that participants did not focus on guessing the purpose of the experiment. On 15% of trials, participants were required to identify the task in which the ball changed colour.

2IFC trials were performed between conditions (*self*_*active*_ v *other*_*observe*_, *self*_*active*_ v *self-* _*observe*_, *self*_*observe*_ v *other*_*observe*_) to calculate bias, and within condition (*self*_*active*_ v *self*_*active*_, *other-* _*observe*_ v *other*_*observe*_, *self*_*observe*_ v *self*_*observe*_) to calculate accuracy. Each participant was paired with a gender-matched partner. To reinforce the social nature of the task, a photo of the participant performing the task accompanied by the word “ active” or “ observe” was shown above the rating bar before each task (Figure 1). To prevent confounds arising from differences in participant performance, participants were told that they were observing their partner’s prerecorded activity, when in fact both *self*_*observe*_ and *other*_*observe*_ conditions were their own prerecorded activity. Participants completed the study over two separate days. The study consisted of five sections: maximum effort estimation, practice trials, prerecording trials, between-condition main experiment, and within-condition main experiment.

#### 2.2.2. Maximum Effort Estimation

The first section, maximum effort estimation, consisted of three sequential 10-second trials with a gravity force that increased exponentially as the ball moved up the ramp. To encourage participants to exert maximum effort a bonus reward of €1 was offered for reaching the top of the ramp (no participant reached the top). The values for each participant’s gravity levels were calculated from these trials by measuring the location of the 80th percentile of the ball’s lateral displacement for each of the three trials, excluding the trial with the lowest value, then averaging the remaining two values and calculating the gravity (*g*_*max*_) opposing the ball at this point. That is, *g*_*max*_ was the maximum gravity level participants could resist for two seconds in two out of three trials. For the experiment, six gravity levels were used, each calculated as a percentage of *g*_*max*_ (Table 1). Participants were informed that there were different levels of difficulty, but were not told how many.

**Table 1.** Gravity Levels as Determined by g_max_

| Gravity level | 1 | 2 | 3 | 4 | 5 | 6 |
| --- | --- | --- | --- | --- | --- | --- |
| % of $g_{max}$ | 20% | 26% | 32% | 38% | 44% | 50% |

#### 2.2.3. Practice Trials

Participants performed six practice trials to become familiar with the task, comparing *self*_*active*_ gravity levels of 1v6, 6v1, 1v4, 5v2, and 3v1. During these trials, participants received feedback on their 2IFC choices to ensure that they understood the paradigm. At no other point during the experiment was feedback given. The gravity differences between the 2IFC tasks (*Δgrav*) for these trials were kept intentionally large to prevent participants from becoming artificially proficient in the *self*_*active*_ condition. Participants could repeat the practice section if they wished.

#### 2.2.4. Prerecorded Trials

Next, participants performed the tasks which were recorded and played back to them during the main experiment as both the *self*_*observe*_ and *other*_*observe*_ conditions. The tasks were presented as 2IFC trials. Participants’ performances were recorded by logging the timing of their correctly ordered alternate left and right key presses. This made it possible to show replays of the participants’ prerecorded tasks within 0.01 seconds of accuracy, without the need for video recording.

To ensure the *observe* tasks matched the fatigue levels of the *active* tasks, we prerecorded tasks for the within-condition main experiment from *self*_*active*_ v *self*_*active*_ trials and tasks for the between-condition main experiment from *self*_*active*_ v *self*_*observe*_ trials. This was based on pilot data showing that back-to-back *self*_*active*_ trials took longer than 2IFC trials with one *active* and one *observe* task. Participants were instructed to take breaks if they felt fatigued during the prerecording stage and were encouraged to move the ball up the ramp as fast as they could. Prerecorded trials were excluded if participants paused for more than 3 seconds between key presses in order to eliminate easily identifiable *observe* tasks where the ball rolled back extensively.

Participants were told they were observing a partner performing the *other*_*observe*_ tasks, but were actually watching their own prerecorded activity. This prevented potential confounds arising from differences in participant ability. To ensure that participants believed they were observing their partner, they had to wait until both they and their partner had finished the prerecording section of the experiment, at which point the experimenter manually transferred files between the two computers using a USB stick. We assessed whether participants believed they were observing a partner’s trials in a post-experiment interview in which they were asked, “ How do you think your partner compared to you in ability? Were they better, worse or the same?” Participants were then informed that they had seen their own prerecorded trials for the *other*_*observe*_ condition and were asked, “ On a scale of 1–10, with 1 being not at all, and 10 being completely, to what degree did you suspect the other participant’s activity wasn’t their own?” If participants indicated any scepticism after the first question, or if they answered the follow-up question with a rating of 10, their data was removed from the analysis. Only one participant was removed due to these considerations.

#### 2.2.5. Main Experiment: Between-Condition 2IFC

The between-condition and within-condition portions of the study were performed on different days, with the order counterbalanced across participants. Participants attended with their partners on the first day of the study, but for logistical reasons could attend the second day separately. To measure self/other and active/observe bias participants performed 2IFC trials comparing *self*_*active*_ v *other*_*observe*_, *self*_*observe*_ v *other*_*observe*_, and *self*_*active*_ v *self*_*observe*_. Each trial was categorised by the difference in gravity level between the two tasks (*Δgrav*). For between-condition trials, *Δgrav* was given by *Δgrav* = *glevel*_*i*_ − *glevel*_*j*_, where *glevel*_*i*_ and *glevel*_*j*_ are the gravity level of conditions *i* and *j*, respectively. Six steps were used: *Δgrav* = −5, −3, −1, +1, +3, +5. These comprised of comparisons of the following gravity levels (italicized): ±1: *1v2, 2v3, 3v4, 4v5, 5v6*; ±3: *1v4, 2v5, 3v6*; ±5: *1v6*. Twenty trials were performed for each *Δgrav* for a total of 120 trials per condition, or 360 trials altogether. Trials were presented in blocks of 30. The blocks were presented in pseudorandomised order: Each set of three blocks comprised one block from each of the three 2IFC comparisons presented in a randomized order. The *Δgrav* values and their associated gravity levels were balanced across blocks and presented in random order.

#### 2.2.6. Main Experiment: Within-Condition 2IFC

Within-condition 2IFCs allowed us to compare participant accuracy in *active* and *observe* conditions. For these trials, *Δgrav* was given by *Δgrav* = |*glevel*_1_ − *glevel*_*2*_|, where *glevel*_1_ and *glevel*_*2*_ are the gravity levels of the first and second task in each trial. Five *Δgrav* levels (1–5) were composed of the following 2IFCs (gravity levels italicised), respectively: 1: *1v2, 2v3, 3v4, 4v5, 5v6*; 2: *1v3, 2v4, 3v5, 4v6*; 3: *1v4, 2v5, 3v6*; 4: *1v5, 2v6*; 5: *1v6*. Each *Δgrav* level was appeared in 20 trials for a total of 100 trials per condition, which were performed in four blocks of 25 trials. These four blocks for each of the three conditions were pseudorandomized as in the between-condition main experiment.

### 2.3. Analysis and Hypotheses

All analysis was performed using MATLAB (2016a, The MathWorks, Inc., Natick, Massachusetts, United States). The following analysis and hypotheses were preregistered at https://osf.io/6pwvf. Exploratory analysis was also performed and is included in a separate section (2.3.5) for clarity.

#### 2.3.1. Hypothesis 1: Bias In self_active_ v other_observe_ Comparisons

We hypothesized that participants would exhibit a bias when comparing the perceived difficulty of tasks they actively performed with tasks they observed their partner performing. This was measured by fitting the 2IFC binary rating data from all *self*_*active*_ v *other*_*observe*_ trials to a psychometric function using the MATLAB function psignfit (Schütt et al., 2016). The *Δgrav* level for each trial provided the stimulus level data (the independent variable) and the dependent variable was set as the proportion of trials rated as easier for *other*_*observe*_. The data was fit using a cumulative Gaussian distribution (options.sigmoidName = ‘norm’), parameters for upper and lower asymptote were set as equal but allowed to vary (options.expType = ‘equalAsymptote’), the threshold was set to detect the pooint of subjective equality (PSE; options.threshPC = .5), and threshold confidence intervals were set to 0.95 (options.confP = .95). All other settings were left as default. The PSE for *self*_*active*_ v *other*_*observe*_ was calculated for each individual. Hypothesis 1 was assessed using a two-tailed one sample t-test with significance set at *p* < 0.05.

#### 2.3.2. Hypothesis 2: Biases For Active/Observe and Self/Other

We hypothesized that two dissociable biases would be detected: one driven by asymmetric sensory information on *active* v *observe* trials, and another driven by attributional differences on *self* v *other* trials. This was predicted to hold even if Hypothesis 1 showed no net bias—in that case, the biases would still be present, but would offset each other.

We dissociated the potential sources of bias—active/observe asymmetry and self/other asymmetry—by calculating the relative PSE between the conditions. The active/observe bias (*PSE*_*active v observe*_) was calculated as the difference in PSE of the *self*_*active*_ v *other*_*observe*_ and the *self*_*observe*_ v *other*_*observe*_ conditions (Equation 1), as this comparison isolated the bias attributed to sensory differences when comparing task difficulty with that of a partner. Similarly, the self/other bias (*PSE*_*self v other*_) was calculated as the difference in PSE between *self*_*active*_ v *other*_*observe*_ and *self*_*active*_ v *self*_*observe*_ (Equation 2). This allowed us to isolate the bias stemming from attributional differences.

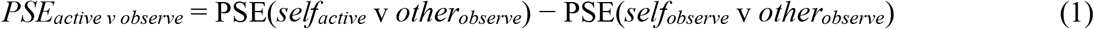

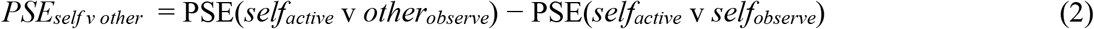

For each participant, PSEs were measured for the three between-condition 2IFC comparisons (*self*_*active*_ v *other*_*observe*_, *self*_*observe*_ v *other*_*observe*_, and *self*_*active*_ v *self*_*observe*_) using the psignfit function with the same settings as Hypothesis 1. The *Δgrav* level for each trial provided the stimulus level data (the independent variable), while the dependent variable was set as the proportion of trials rated as easier for Condition 2 in each comparison. Conditions 1 and 2 were defined as shown in Table 2.

**Table 2.** Definitions of Conditions 1 and 2 for Psychometric Between-Condition Analysis

| 2IFC Comparison | Condition 1 | Condition 2 |
| --- | --- | --- |
| *self*<sub>active</sub> v *other*<sub>observe</sub> | *self*<sub>active</sub> | *other*<sub>observe</sub> |
| *self*<sub>observe</sub> v *other*<sub>observe</sub> | *self*<sub>observe</sub> | *other*<sub>observe</sub> |
| *self*<sub>active</sub> v *self*<sub>observe</sub> | *self*<sub>active</sub> | *self*<sub>observe</sub> |

*Note.* As the main analysis of interest was *self<sub>active</sub>* v *other<sub>observe</sub>*, *self<sub>active</sub>* was set as Condition 1 and *other<sub>observe</sub>* was set as Condition 2. Because *self<sub>observe</sub>* was used as a control condition to extract the two biases of interest (*self* v *other*, *active* v *observe*), it replaced *self<sub>active</sub>* in the second comparison and *other<sub>observe</sub>* in the third comparison.

The presence of bias for active/observe and self/observe was analysed using a linear mixed effects model (MATLAB function fitlme; Pinheiro & Bates, 1996). Values for *PSE(self*_*active*_ v *other*_*observe*_*), PSE(self*_*active*_ v *self*_*observe*_), and *PSE(self*_*observe*_ v *other*_*observe*_) were entered as dependent variables, with categorical parameters indicating whether the 2IFC involved a self/other or active/observe asymmetry added as fixed effects parameters. Random effects were added to allow for random intercepts for each participant as well as for the two sources of bias. To test Hypothesis 2, the full linear mixed effects model was compared to reduced models with fixed effects for only s*elf* v *other, active* v *observe* or neither (*null*) using likelihood ratio testing (MATLAB function compare(lme,altlme); Hox & Maas, 2005). Hypothesis 2 was also analysed using a two-tailed one sample t-test (α = 0.05) to determine whether *PSE*_*self v other*_ and *PSE*_*active v observe*_ were significantly different from zero.

#### 2.3.3. Hypothesis 3: Greater Accuracy in Active tasks relative to Observe tasks

We hypothesized that participants would be more accurate in their 2IFC ratings when they had performed the effortful task than when they had observed it. To assess this, we fit binary ratings from the within-condition 2IFC trials of the three conditions (*self*_*active*_, *other*_*observe*_, and *self*_*observe*_) with a psychometric function. A threshold we refer to here as the just-noticeable difference^1^ (JND) was set to detect the point that participants correctly identify the more difficult task in 80% of trials (after adjustment to account for lapse rates as per Schütt et al., 2016). This threshold was based on pilot studies, and set to 80% rather than the more standard 75% to account for some participants achieving higher accuracy for the most difficult comparisons (Δ*grav* = 1).

The 2IFC data was fit with the psignfit function (Schütt et al., 2016) using a cumulative Gaussian distribution (options.sigmoidName = ‘norm’) and the parameter for lower asymptote was locked at 0.5 while the upper asymptote was allowed to vary (options.expType = ‘2AFC’).

The threshold was set at 0.8 (options.threshPC = .8). The stimulus level data was provided by the Δ*grav* value, while the dependent variable was the proportion of trials correctly rated at each Δ*grav*.

Statistical comparison of *self*_*active*_, *other*_*observe*_, and *self*_*observe*_ was performed using a one-way analysis of variance (ANOVA). Multiple comparisons with a Bonferroni correction was used to provide statistical tests of significance between each of the three conditions. To assess Hypothesis 3 and determine the role of sensory availability differences in *active* and *observe* trials, a repeated-measures ANOVA was performed with the independent variables of *active/observe* and *self/other*. Statistical significance was set at *p* < 0.05 for both tests.

#### 2.3.4. Hypothesis 4: Do Differences in Accuracy Correlate With Bias

We hypothesized that the level of active/observe bias would correlate with greater accuracy differences between *active* and *observe* trials, but the self/other bias would not correlate with accuracy differences between *self* and *other* trials. Pilot data suggested that bias between *active* and *observe* tasks may be driven by asymmetries in accuracy when either performing or observing a task. To explore this, a Pearson’s correlation (MATLAB function corrcoef) was performed between participants’ *PSE*_*active v observe*_ and the JND between the *active* and *observe* conditions, as given by Equation 3, where *PSE*_*active v observe*_ is the bias between *active* and *observe* trials (see 2.3.2. Hypothesis ***2***) and the right side of the equation captures the accuracy difference between the *observe* within-condition trials and the *active* within-condition trials.

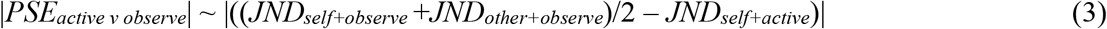

The absolute value of both sides of the equation is taken under the assumption that the accuracy difference, but not the direction of the difference, drives the magnitude of the active/observe bias. The statistical threshold for significance was set at *p* < 0.05.

The same analysis was also performed for *PSE*_*self v other*_ with the expectation that this relationship would not be correlated, as participants would not receive any information which would increase their accuracy in either the *self* or *other* conditions.

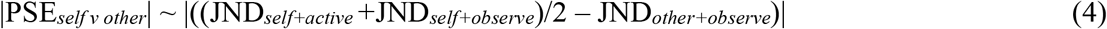

#### 2.3.5. Exploratory Analysis

The preregistered analysis focused solely on the binary 2IFC decision data. However, continuous rating data was also captured for each trial by measuring the displacement of the ratings bar. Therefore, to complement the psychometric modelling of the preregistered analysis, the analysis of Hypotheses 1 and 2 was repeated using the continuous rating data from the between-condition comparisons. Using the continuous rating data made it possible to capture subtleties in perceptual judgement that the binary 2IFC data may miss.

We also used the continuous rating data to analyse the effect of task difficulty on active/observe bias. If bias in social comparison was driven by sensory asymmetries, the perceived bias would be expected to increase as task difficulty increased. To test this, bias was calculated for each condition for trials where multiple combinations of gravity levels were used to form the same *Δgrav* (i.e., trials where Δ*grav* = +1, −1, +3, and −3). As Δ*grav* = ±5 was only possible in one combination (6 v 1 and 1 v 6), it were not used in this analysis. For each permutation, we tested active/observe and self/other bias for significant deviation from zero using two-sided one sample t-tests. Furthermore, a repeated measures ANOVA assessed whether there was a significant interaction between relative task difficulty and active/observe bias for each of the four *Δgrav* levels by comparing the participant ratings with an independent variable for the gravity level for Condition 1 and a categorical independent variable for the presence or absence of active/observe asymmetry. Significant effects in ANOVA were followed up by post hoc pair-wise comparisons (MATLAB function multcompare, with Bonferroni correction).

Finally, based on results from this exploratory analysis of task difficulty, a linear mixed effects model was run using the continuous rating data from all trials in order to determine whether task difficulty was driving the level of active/observe bias. The model included fixed effects parameters indicating whether the 2IFC trial involved a self/other asymmetry, an active/observe asymmetry, the Δ*grav* for each trial, the condition presented first (to capture variance in how participants were influenced by presentation order), and the gravity level of Condition 1 for each trial (to capture the effect of overall task difficulty). As task difficulty was expected to drive the amount of bias ascribed to the active/observe asymmetry, the fixed effects parameter for the gravity level of Condition 1 was included as an interaction term with the parameter for active/observe asymmetry. The gravity level of Condition 1 was also included as an interaction term with the parameter indicating which condition was presented first. As random effects, we included random intercepts for each participant, and by-participant random slopes for each fixed effect. We performed nested model comparisons with reduced versions of this full model to evaluate model performance. As with all previous model comparisons, ΔBIC (Bayesian information criteria; Schwarz, 1978) and ΔAIC (Akaike’s information criteria; Burnham & Anderson, 1998) were performed; a theoretical likelihood test (Hox & Maas, 2005) was used in the event of disagreement between these two criteria.

## 3. Results

### 3.1: Hypothesis 1: No Bias Between *self*_*active*_ and *other*_*observe*_

No bias between *self*_*active*_ and *other*_*observe*_ was detected at the group level (*M* = 0.106, *SD* = 0.75, *t*(50) = −1.01, *p* = 0.3166). Participants exhibited PSEs that ranged from underestimating (Figure 2A, positive values) to overestimating (Figure 2A, negative values) the difficulty of tasks they performed themselves. However, as there was no systematic bias across all participants, we were unable to reject the null hypothesis. Analysis using the continuous rating data also found no significant bias (*M* = −0.1066, *SD* = 0.5732, *t*(50) = −1.33, *p* = 0.1904) (Supplementary Figure S1A).

**Figure 2.**
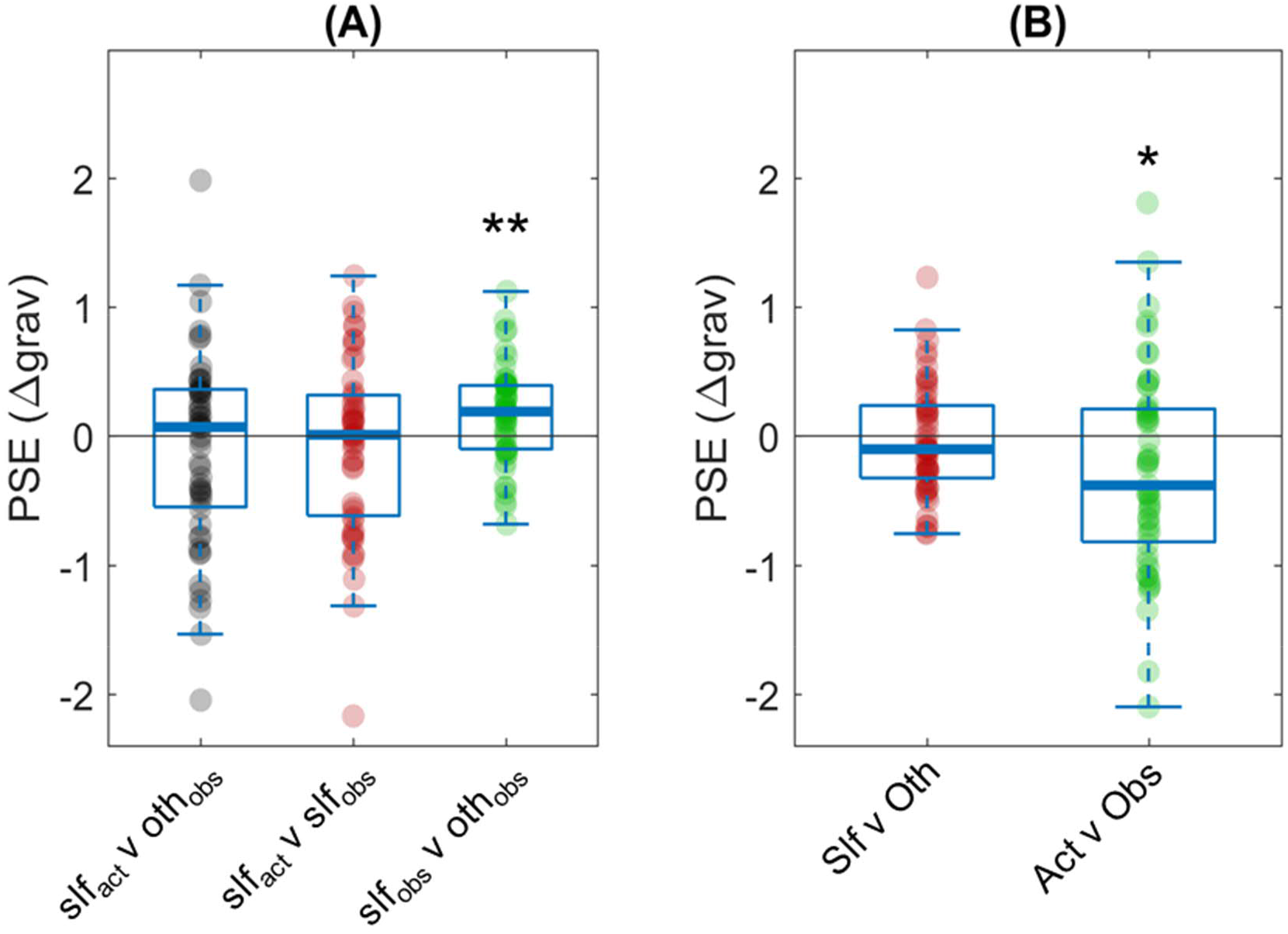
Point of Subjective Equality (PSE) Between Conditions. *Note*. Both panels plot the PSE between conditions as calculated by psychometric functions, with negative values indicating an overestimation of the difficulty of the condition listed first (i.e., Condition 1). Panel A: Two-interval forced choice results between the three conditions: performing the task (slf_act_), watching a partner perform the task (oth_obs_), and watching a recording of oneself performing the task (slf_obs_). Panel B: Attribution bias (Slf v Oth) and sensory asymmetry bias (Act v Obs) for all participants. \**p* ≤ 0.05, \*\**p* ≤ 0.01.

It is worth noting that no bias was detected in the *self*_*active*_ v *self*_*observe*_ condition (*M* = −0.0797, *SD* = 0.68, *t*(50) = −0.841, *p* = 0.4044; Figure 2A). However, participants significantly underestimated the difficulty of their own trials in *self*_*observe*_ v *other*_*observe*_ comparisons (*M* = 0.168, *SD* = 0.3762, *t*(50) = 3.19, *p* = 0.0024), even though these trials compared identical stimuli. This result may indicate evidence of metacognitive processes, where participants consciously accounted for a potential egocentric bias due to the comparative nature of the task (we address this possibility further in the Discussion section).

### 3.2 Hypothesis 2: Active/Observe Bias Detected, But No Bias for Self/Other

Participants exhibited no self/other bias as a group (*M* = −0.0267, *SD* = 0.4366, *t*(50) = −0.4360, *p* = 0.6647, *d* = −0.037; Figure 2). However, a significant active/observe bias emerged, with participants overestimating the difficulty of tasks they performed themselves (*M* = −0. 2746, *SD* = 0.7859, *t*(50) = −2.746, *p* = 0.0159, *d* = −0.462). Hypothesis 2 stated that both biases would be detected; the null hypothesis therefore cannot be rejected. Instead, these results point toward a single source of bias for participants in this paradigm, driven by the asymmetries of performing or observing the task.

The mixed effects modelling mirrored this finding, with AIC and BIC model selection choosing the Active v Observe and Null model, respectively (Table 3), while a likelihood ratio test suggested that the Active v Observe model explained the data significantly better (*χ*^2^(1) = 4.4984, *p* = 0.0332). In the Active v Observe model, performing the task increased perceived difficulty by 0.261 ± 0.082 (standard errors) gravity levels. However, in the analysis using continuous ratings, both AIC and BIC model comparisons selected the Active v Observe model. This finding was further supported by a likelihood ratio test: *χ*^2^(1) = 46.15, *p* <0.0001 (Supplementary Table S1).

**Table 3.** Attribution Bias and Sensory Asymmetry Bias Model Comparison

| Model | ΔAIC | ΔBIC | Log Likelihood | R<sup>2</sup> (adjusted) |
| --- | --- | --- | --- | --- |
| Full | 1.92 | 5.44 | -132.41 | 0.3789 |
| Active v Observe | 0 | 0.49 | -132.45 | 0.3823 |
| Self v Other | 4.53 | 5.02 | -134.72 | 0.3714 |
| Null | 2.53 | 0 | -134.72 | 0.3758 |

*Note.* Mixed-effect models featuring fixed effects for both attribution bias and sensory asymmetry bias (Full) or only one bias (Self v Other, Active v Observe) were fit with the between-condition point of subjective equality (PSE) data. Akaike's information criteria (AIC) and Bayesian information criteria (BIC) differed in their assessment of the winning model between PSE<sub>active</sub> and the null model indicating no bias. A likelihood ratio test assessed that the PSE<sub>active</sub> model significantly outperformed the Null model ( $\chi^2(1) = 4.50$ , $p = 0.033$ ). Gray backgrounds denote winning models.

**Table 4.** Repeated Measures ANOVA Results for Task Difficulty and Active/Observe Bias

| $\Delta grav$ | IV | DF | F | p | $\eta_p^2$ |
| --- | --- | --- | --- | --- | --- |
| +1 | gl1 | (1,50) | 17.65 | 0.0001 | 0.261 |
|  | Active | (1,50) | 19.45 | <0.0001 | 0.280 |
|  | gl1*Active | (1,50) | 24.26 | <0.0001 | 0.327 |
| +3 | gl1 | (1,50) | 4.31 | 0.043 | 0.079 |
|  | Active | (1,50) | 27.34 | <0.0001 | 0.354 |
|  | gl1*Active | (1,50) | 26.45 | <0.0001 | 0.346 |
| -1 | gl1 | (1,50) | 3.88 | 0.054 | 0.072 |
|  | Active | (1,50) | 0.01 | 0.915 | 0.000 |
|  | gl1*Active | (1,50) | 0.04 | 0.568 | 0.007 |
| -3 | gl1 | (1,50) | 2.53 | 0.118 | 0.048 |
|  | Active | (1,50) | 0.05 | 0.823 | 0.001 |
|  | gl1*Active | (1,50) | 0.14 | 0.713 | 0.003 |

*Note.* Two independent variables (IV), *gl1*—the difficulty level of the Condition 1 task—and *Active*—a categorical parameter for whether there was an active/observe asymmetry—were assessed against group bias for the four $\Delta grav$ – gravity level 1 – gravity level 2.

### 3.3. Hypothesis 3: Greater Accuracy in Active Tasks Than Observe Tasks

We next compared the accuracies (JNDs) between the three experimental conditions (Figure 3). Participants were more accurate in *self*_*active*_ trials (*M* = 2.29 ± 0.98) than in *self*_*observe*_ (*M* = 3.11 ± 1.20) and *other*_*observe*_ (*M* = 2.86 ± 1.01) trials. A repeated measures ANOVA revealed a statistically significant difference between conditions (F(2,100) = 19.791, *p* <10^−7^). Post hoc tests showed a significant difference between *self*_*active*_ and *self*_*observe*_ (*p* = 0.0003) and between *self*_*active*_ and *other*_*observe*_ (*p =* 0.019), but not between the two *observe* conditions (*p* = 0.698). Based on these results, we were able to reject the null hypothesis.

**Figure 3.**
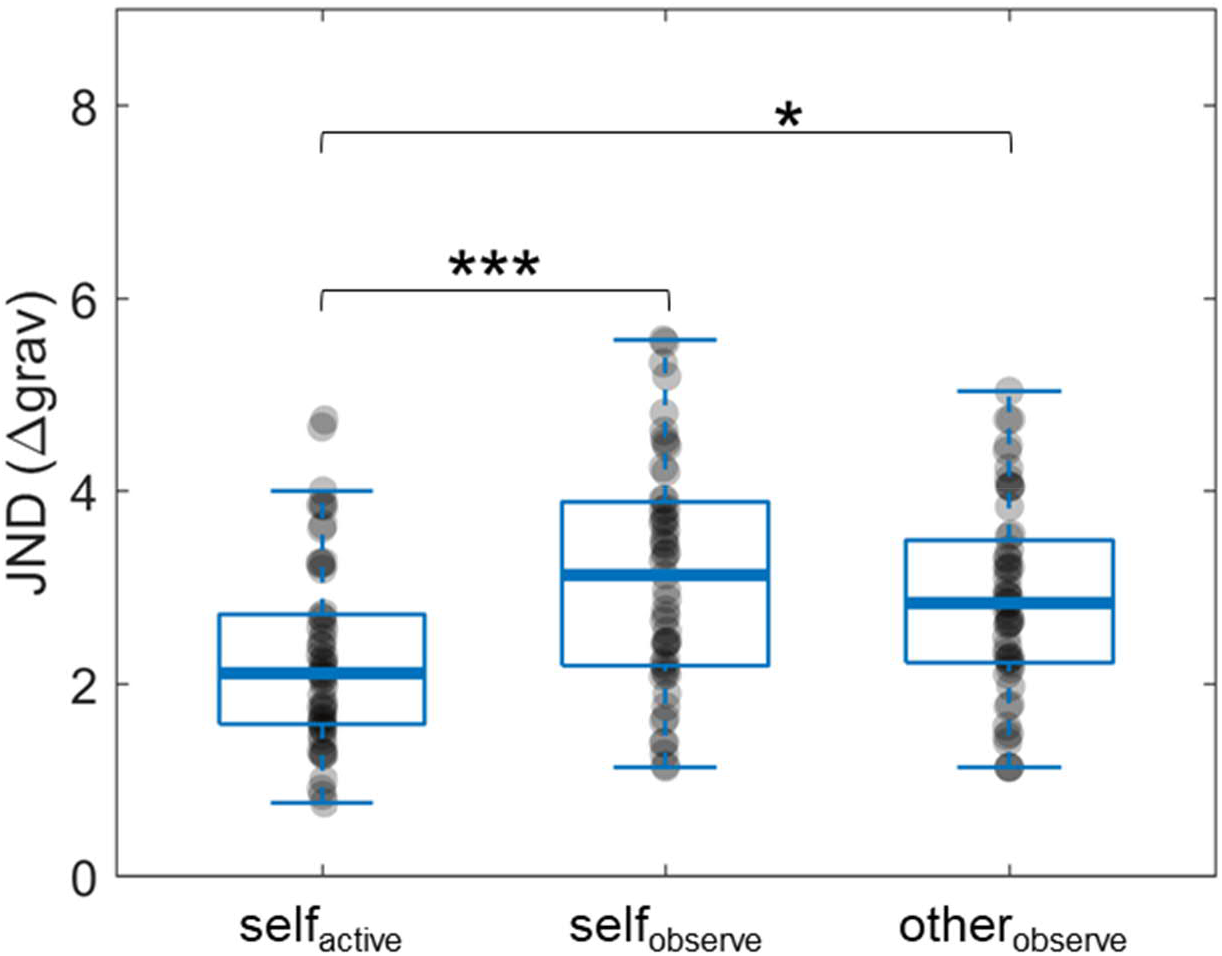
Within-Condition Just-Noticeable Difference (JND) Results. *Note*. Accuracy was measured by JND, defined here as the difference in gravity levels (Δ*grav*) where participants correctly identified the more difficult task in 80% of trials. Smaller JND indicates greater accuracy. \**p* < 0.05, \*\*\**p* < 0.001.

### 3.4. Hypothesis 4: No Relationship Between Accuracy and Bias

No correlation was found between *PSE*_*active v observe*_ and active/observe accuracy differences using either the preregistered (Pearson’s *r* = 0.08, *p* = 0.57) or the exploratory (*r* = 0.16, *p* = 0.27) model of accuracy driven bias (Supplementary Figures S2, S3). The same analysis for self/other also found no significant correlation (preregistered: *r* = −0.0008, *p* = .996; exploratory: *r* = −0.0016, *p* = .991). We found no evidence for a relationship between accuracy differences in the different task conditions and bias.

### 3.5. Exploratory Results

Our *a priori* hypothesis, that bias is driven by accuracy differences between active/observe, was not supported. We therefore explored an alternative explanation, that bias was driven by sensory asymmetries between *active* and *observe* conditions. If this was correct, we would expect bias to increase with increased task difficulty. We used the continuous rating data to analyse and model the effect of task difficulty (Δ*grav*) on active/observe bias: We performed a repeated measures ANOVA with a predictor for Condition 1 gravity level (*gl1*) and a categorical predictor for the presence of active/observe asymmetry, as well as an interaction between these two predictors. Only the between-condition trials where unique permutations for each Δ*grav* levels occurred (Δ*grav* = +3, +1, −1, −3) were included in the ANOVA (see also Methods section 2.3.5. We found that as the overall task difficulty increased, participants exhibited greater bias overestimating the difficulty of *active* task relative to *observe* trials (Figure 4)— albeit only when the *active* task was more difficult than the observe task (Δ*grav* = +1, +3).

**Figure 4.**
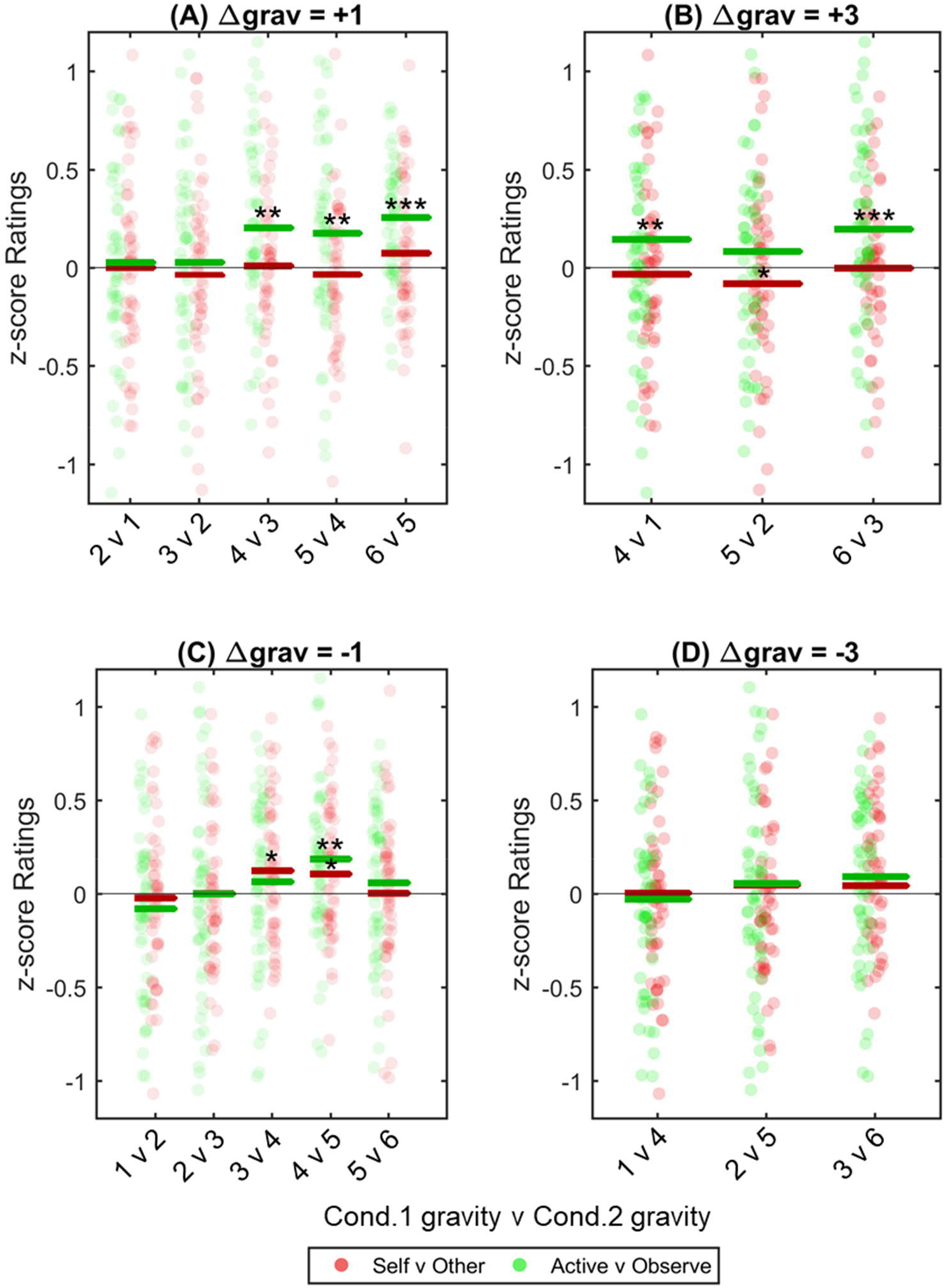
Effect of Task Difficulty on Bias. *Note*. Biases for each participant were calculated for all permutations for *Δgrav* = 3, 1, −1, −3. These were pooled across all participants and graphed to show bias as task difficulty increased. Panels A, B: Post hoc one sample t-tests show that when participants actively performed the more difficult task there was a significant bias for all trials where Condition (Cond.) 1 was ≥ gravity level 4 (apart from 5v2). Panels C, D: This effect was not seen when participants performed the easier of the two tasks. Data was analysed with a repeated measures two-way ANOVA, with an interaction between bias type and task difficulty significantly affecting the variance and with post hoc one sample t-tests applied to each gravity level permutation. *\*p* < 0.05, **p < 0.01, \*\*\**p* < 0.001.

Finally, we reran our mixed-effect modelling using the continuous rating data with an added fixed effect for overall task difficulty, as indicated by the gravity level of Condition 1 in each trial. The gravity level of Condition 1 was added as an interaction term with the fixed effect for active/observe asymmetry (and as an interaction term with the fixed effect for task presentation order). Both BIC and AIC analysis selected the Full model as the best fit for the data (**Table 5**).

**Table 5.** Exploratory Model Comparison Assessing the Interaction of Task Difficulty and Bias

| Model | $\Delta AIC$ | $\Delta BIC$ | $R^2$ |
| --- | --- | --- | --- |
| Active + Self + w | 118.99 | 56.44 | 0.600 |
| Active + w | 117.27 | 46.91 | 0.600 |
| Self + w | 129.36 | 59.00 | 0.600 |
| Null | 127.60 | 49.42 | 0.600 |
| Full | 0.00 | 0.00 | 0.603 |
| Active + g * w | 46.12 | 38.30 | 0.602 |
| Active * g + w | 68.44 | 60.62 | 0.602 |
| Active + g + w | 114.37 | 98.74 | 0.601 |
| g + w | 124.70 | 101.25 | 0.601 |

*Note.* Using the subjective rating data, the preregistered models from Hypothesis 2—with categorical parameters for active/observe asymmetry (Active) or self/other asymmetry (Self)—were compared with models incorporating the gravity level of Condition 1 ( $g$ ) as an interaction factor. The Full model included $g$ as an interaction factor with both active/observe asymmetry and task presentation order. Adding this factor significantly improved the model fit according to both AIC and BIC assessment. All models contained a parameter that captured variance caused by task presentation order ( $w$ ) which interacted with $g$ in the Full model (Full model ∼ active \* $g$ + $g$ \* $w$ , Null model $\sim w$ ).

## 4. Discussion

This study examined the effect of attribution (self/other) and availability (active/observe) asymmetries on social comparison of effort. Including a condition where participants observed their own prerecorded activity allowed us to dissociate between the effect of self/other and active/observe asymmetries on egocentric bias. Our results showed that only active/observe asymmetries contributed to egocentric bias; self/other asymmetries had no significant effect. The bias measured in this study was small, with the psychometric modelling using the binary data indicating a mean bias of ∼0.27 gravity levels, or 1.6% of a participants’ maximum effort as measured by *g*_*max*_ (the exploratory analysis using continuous data estimated a more substantial ∼0.6 gravity levels, or 3.6% of *g*_*max*_). However, this small effect size was expected, and was one of the primary motivations behind preregistering the study.

Our study provides insights into the underlying mechanism that drives egocentric bias. We predicted that greater access to sensory information while performing a task would lead to greater accuracy relative to simply observing a task, and that differences in accuracy between these two states would correlate with the degree of bias exhibited. Our hypothesis was not supported by the results: While participants were significantly more accurate when comparing *active* tasks than *observe* tasks, there was no evidence of a correlation between differences in *active* and *observe* accuracy and the amount of bias.

However, our exploratory analysis did support a correlation between overall task difficulty and active/observe bias, albeit only in tasks where participants performed, not observed, the more difficult task.

It is worth noting that the task difficulty in this study was intentionally relatively low. The study took an average of 4 hours to complete; therefore, to avoid excessive discomfort or participant attrition, the task was set to a level around 60% of that used in previous iterations of the ball and ramp paradigm (Pooresmaeili et al., 2015; Rollwage et al., 2020). Given our results correlating increased bias with task difficulty, future studies incorporating greater task difficulty may find greater active/observe bias.

We found no bias correlating with self/other asymmetries. However, this finding may be a direct result of the experiment’s incentive payments. Participants were explicitly rewarded for accuracy, which may have motivated them to override any overt or motivated biases. While explicitly rewarding accuracy was intentional—we wanted to detect inherent egocentric bias rather than motivated bias—it is possible that our expectation of a self/other bias was thwarted by our incentive structure. This possibility was supported by results suggesting participants were actively deploying metacognitive strategies to compensate for any inherent self/other bias. This was particularly evident in the *self*_*observe*_ v *other*_*observe*_ condition, where participants compared identical stimuli and yet rated the tasks attributed to their partner as significantly more difficult (Figure 1A). It is important to note, however, that this bias in the *self*_*observe*_ v *other*_*observe*_ condition did not affect the study’s ability to extract the unmotivated self/other bias, which was captured by the comparison between the main condition (*self*_*active*_ v *other*_*observe*_) and the relevant control (*self*_*active*_ v *self*_*observe*_).

Metacognitive strategy may also explain the exploratory result that found that task difficulty was correlated with greater bias only in trials where participants performed the more difficult task themselves (Figure 4). A participant countering their own egocentric bias to maximise their reward would focus on the correctness of their binary rating but not the accuracy of their continuous rating. This could lead participants to strategically apply a counterbias to trials that fell below a certain threshold of perceptual certainty. Given that our data indicated the participants overestimated *active* tasks, participants would, on this account, have greater perceptual certainty on trials where the *active* task was more difficult, while tasks where the *observe* task was more difficult would be perceived with less certainty and therefore be more likely to require metacognitive intervention. Further studies (e.g., using graded wagering; Moreira et al., 2018) are required to test this proposed mechanism.

This study focused solely on bias inherent to the egocentric perspective. As such, a range of potential biases driven by self/other asymmetries were not addressed. Well documented biases that are better described by this asymmetry (e.g., the endowment effect; Kahneman et al., 1991) were necessarily excluded by our paradigm. Furthermore, the social aspect of our experiment paired strangers from relatively homogenous demographics, and as such did not explore potential attributional biases produced by perceived differences between oneself and others (Ashkanasy, 1997; Green & McClearn, 2010). Therefore, this study makes no comment on the existence or prevalence of self/other biases in general, but rather aims to better characterise the bias inherent in egocentric perception.

## 5. Conclusions

Our study featured a novel paradigm that allowed us to dissociate between egocentric bias driven by active/observe and by self/other asymmetries. We found that participants perceived tasks they performed to be harder than tasks they observed, but found no bias correlating with whether the participant or their partner performed the task (*self* or *other*). Furthermore, exploratory analysis suggested that as the effortful task became more difficult, participants exhibited greater bias towards overestimating *active* tasks.

This study contributes to the field of egocentric biases by dissociating two potential sources of bias (self/other and active/observe asymmetries) using a control condition where participants observe their own prerecorded activity. This makes it possible to separate two proposed biases that are often understood as synonymous within the literature, and suggests that at least some biases previously attributed to self/other asymmetries may be better explained by active/observe asymmetries.

## Acknowledgements

This work was supported by a seed fund grant from Leibniz ScienceCampus *Primate Cognition*, Göttingen, Germany to IK and AP. AP was supported by an ERC Starting Grant (716846, *Rewarded Perception*). Deborah Ain reviewed and edited the manuscript.

## Authors’ contributions

**Caedyn Stinson:** Conceptualization, Methodology, Software, Formal Analysis, Investigation, Resources, Data Curation, Writing, Visualisation, Project administration **Arezoo Pooresmaeili:** Conceptualization, Methodology, Software, Formal Analysis, Resources, Writing – Review & Editing, Supervision, Funding Administration **Igor Kagan:** Writing – Review & Editing, Funding Administration.

## Data availability

The datasets generated and analysed in the current study are available from the corresponding author upon request, and will be uploaded to OSF (https://osf.io/35qz2/?view_only=9c61d0374cf64c889b6f45a252b51653).

## Footnotes

1 Typically, the JND measures an initial detection threshold. We are appropriating the term to denote a measure for noticeable differences between gravity levels of two tasks.

