## Supplementary Material for "Egocentric Bias in Effort Comparison Tasks Is Driven by Sensory Asymmetries, Not Attribution Bias"

**Figure S1**

*Subjective Rating Based Point of Subjective Equality ($PSE_{ratings}$) Analysis of Bias*

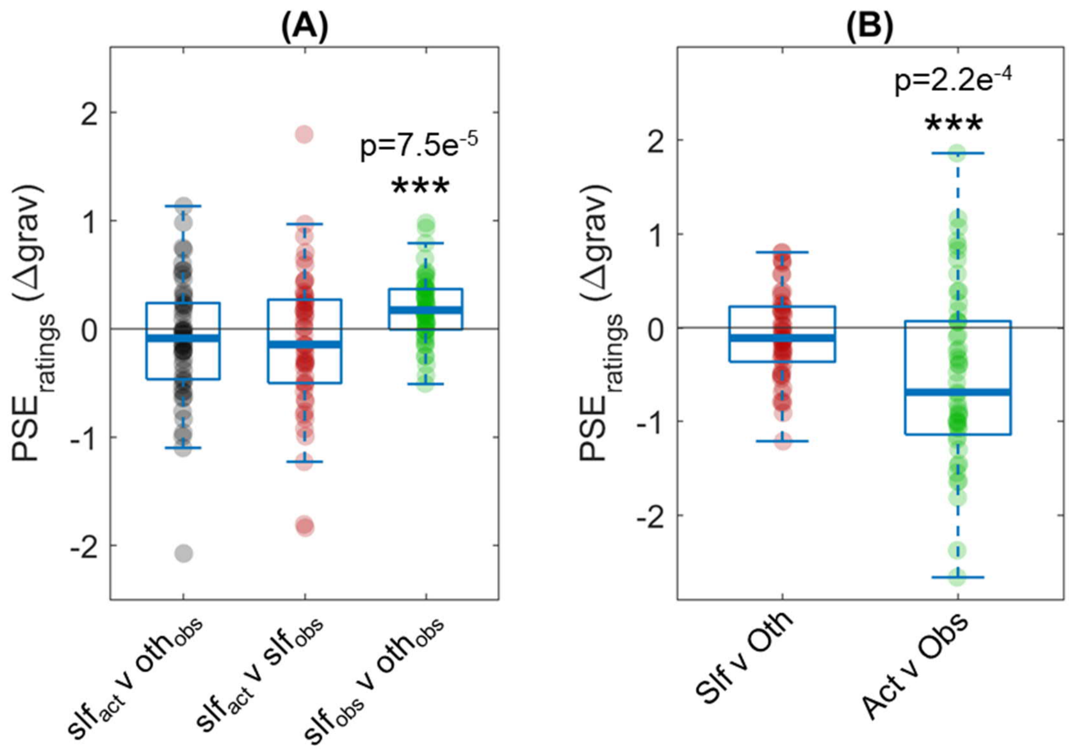


*Note*. Both figures plot the $PSE_{ratings}$, with negative values indicating an overestimation of the difficulty of the condition listed first (i.e. Condition 1). (A) shows biases in ratings between the three conditions, with no bias detected at a group level for $slf_{act}$ v $oth_{obs}$ ($t(50) = -1.33$, $p=0.190$) or $slf_{act}$ v $slf_{obs}$ ($t(50) = -1.33$, $p = 0.191$). Participants significantly underestimated the difficulty of their own trials in $slf_{obs}$ v $oth_{obs}$ comparisons ($t(50) = 4.31$, $p <0.0001$). (B) displays attribution bias (Slf v Oth) and sensory asymmetry bias (Act v Obs) for all participants. At a group level, only sensory asymmetry bias was detected ($t(50) = -3.99$, $p = 0.0002$) with no differences found for attribution bias ($t(50) = - 1.21$, $p = 0.234$).

**Table S1**

*Attribution Bias and Sensory Asymmetry Bias Model Comparison with Rating Data*

| Model | ΔAIC | ΔBIC | Log Likelihood | $R^2$ (adjusted) |
|---|---|---|---|---|
| $PSE_{ratings}$ full | 1.72 | 9.54 | -18041 | 0.3789 |
| $PSE_{ratings}$ active | 0 | 0 | -18042 | 0.3823 |
| $PSE_{ratings}$ self | 37.51 | 37.51 | -18060 | 0.3714 |
| $PSE_{ratings}$ null | 44.15 | 36.34 | -18065 | 0.3758 |

*Note.* Models featuring both attribution bias and sensory asymmetry bias ($PSE_{ratings}$ full) or only one bias ($PSE_{ratings}$ self and $PSE_{ratings}$ active) were fit with the between-condition PSE data. In contrast to the PSE results, model comparisons using both AIC and BIC were in agreement that the model with a lone bias parameter for *Active v Observe* best explained the data.

**Figure S2**

*Analysis of the correlation between relative accuracy and bias (pre-registered analysis)*

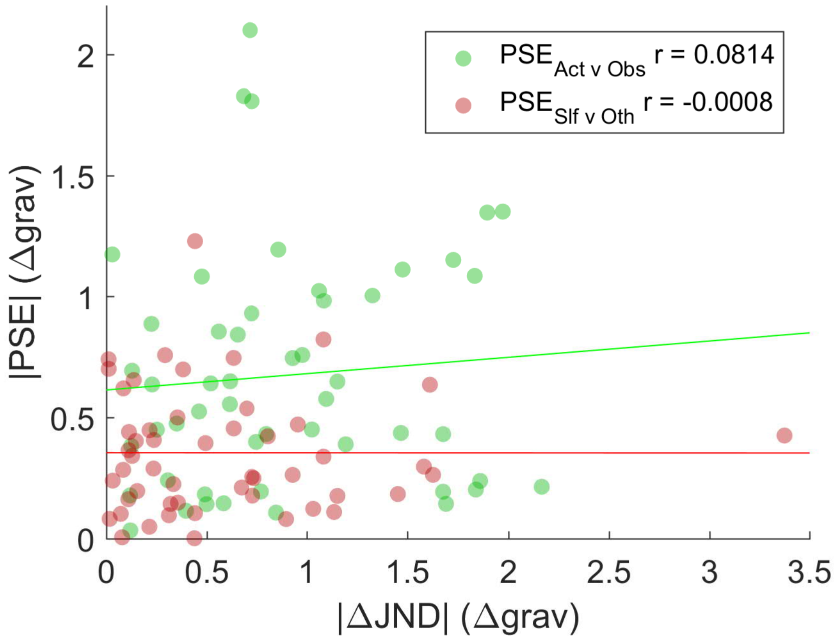


*Note.* The absolute value of participant's bias (|PSE|) is plotted against the differences in their accuracy between either *Active* and *Observe* tasks (green) or *Self* and *Other* tasks (red). The correlation is given by *Pearson's r*, with neither correlation significant (*Active v Observe p* = 0.5702, *Self v Other p* = 0.9958).

**Figure S3**

*Analysis of the correlation between relative accuracy and bias (exploratory analysis)*

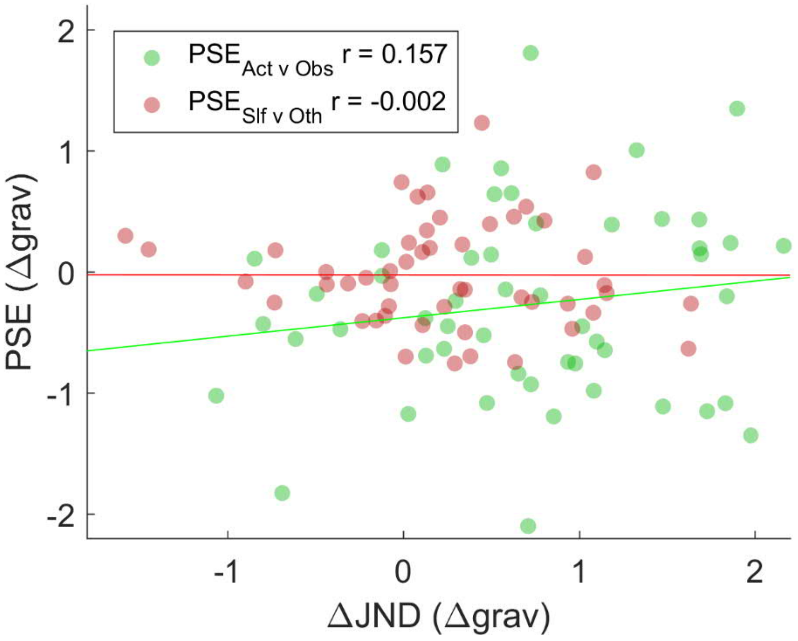


*Note.* The absolute value of participant's bias (|PSE|) is plotted against the differences in their accuracy between either *Active* and *Observe* tasks (green) or *Self* and *Other* tasks (red). The correlation is given by *Pearson's r*, with neither correlation significant (*Active v Observe p* = 0.2706, *Self v Other p* = 0.9914).
